# Cell-to-cell Mathematical modeling of arrhythmia phenomena in excitable media

**DOI:** 10.1101/624247

**Authors:** Gabriel López Garza

## Abstract

In this document are modeled arrhythmias with cellular automaton and ordinary differential equations systems. With an aperiodic, self-similar distribution of two-dimensional arrangement of cells, it is possible to simulate such phenomena as fibrillation, fluttering and a sequence of fibrillation-fluttering. The topology of the cytoarchitecture of a network of cells may determine the initiation and development of arrhythmias.

## 1. Introduction

Following to Wiener and Rosenblueth [33], for mathematical simplicity, we define only two types of arrhythmia in excitable media. One type is known as *Fluttering* is related to reentrant waves of excitation which remain in a self-perpetuating steady state. The second and more complex type of arrhythmia considered in this article is known as *Fibrillation*. Meanwhile fluttering is adequately described employing continuous or cell-to-cell modeling, the fibrillation phenomenon is more difficult to simulate with deterministic models in both partial differential equations and ordinary differential equations. Some authors, [33], [6] consider that fibrillation can only be approached mathematically on a statistical basis, mainly due to the random distribution of anastomosis fibers in the heart. However, in this article, we present a deterministic model in which some complexity is established in the geometry of the cellular network to provide a specific behavior that could be identified to some extent to fibrillation.

One of the thesis of Wiener and Rosenblueth, as the authors state, is that contrary to the hypothesis of Lewis [21], the mechanism of auricular flutter and fibrillation is not a single one. Wiener and Rosenblueth claim that fibrillation may be due to random circus movements. Posteriorly, the authors utilize automaton to explain flutter due to fixed obstacles, such as the Vena Cava. Is worth to mention that in their pioneering work, Automata were purely theoretical. In consequence, the authors did not use computers at all, but only geometric arguments which lead them to right predictions to some extent. On the other hand, to study fibrillation according to their thesis, they introduced the Wiener and Wintner theory of statistical distributions [34]. However, in the case of fibrillation, Wiener and Rosenblueth do not arrive at anything conclusive, since they affirm that solving the obtained equations “is a question for a more complex and difficult mathematical work” a work that, in the opinion of the author, it was never achieved. Nowadays there are articles that attempt to model more complex arrangements in cellular systems [20], but computational constraints strongly determine the possibility of more realistic models. This is one of the reasons that explain why cellular automaton are still in use.

Many years had passed since the publication of Wiener and Rosenblueth and the present paper, and yet, the primary mechanisms of flutter and fibrillation are not fully understood. Researchers still have incomplete knowledge of how arrhythmias such as ventricular fibrillation begin and develop. However, the paradigmatic Hodgkin-Huxley mathematical model of excitable cells discovered in the sixties had provided researchers with an indispensable tool. Also, of course, the increasing power of computers make possible the incorporation of a high number of observables in the modeling of individual cells. Moreover, nowadays some of the randomnesses of arrhythmias may be treated into the realm of Complexity Science. Meanwhile, Medical Science evolved, and the classifications of arrhythmias are by far much more sophisticated than those in the forties [29], [37], [28].

In this article, we study some of the consequences obtained by modeling weakly connected networks through different distributions of excitable cells within the network, what we call *the geometry of the network*. In this context, we argue in the 3.1 section why cell-to-cell modeling fits better than the continuous model, at least to model the arrhythmia. We will illustrate in section 2, that neither the diffusivity provided by partial differential equations nor by the cell-to-cell coupling requires a complex dynamics in the cells to produce fibrillation and flutter phenomena. Diffusivity in continuous mathematical modeling is given by elliptic-type operators and in cell-to-cell modeling using *weakly coupled variables* (see section 3.1.1). Nevertheless, we show *in silico* that fibrillation and fluttering can be modeled even by using the simplest cellular automaton and excitable cell models including only few variables.

The main difference between flutter and fibrillation, according to the classic definitions [33], is the randomness of fibrillation as opposed to the regularity of flutter. Randomness precludes sharp, defined wavefronts. The principal contribution of the present work is to introduce some degree of complexity in the net of connections instead of randomness to present an *in silico* phenomena which can be identified with fibrillation. Another novelty in the present paper is that, contrary to the commonly established, fluttering can be produced at a cellular level by a dynamic obstacle formed with a few cells as well as that produced by fixed non-dynamical obstacles (for a definition see section 2.2.1).

## 2. Methods

### 2.1. Individual Cell models

In this work, we use cellular automaton and two-variable models of excitable cells. The primary interest in this article is to study the diffusivity on excitable media and not in modeling dynamics of individual cells. The models included here, as usual, go through four stages [35]: resting, exciting, excited, and refractory states; also, the models of coupled cells provide solitary waves flexible enough to flutter and fibrillate. In this way, the models represent observables in real tissue to some extent (for a mathematical definition of *observable* see section 3.1.1). The convenience and relevance of utilizing more complex models of individual cells is discussed in sections 5, and 6.

#### 2.1.1. Cellular automaton model

The cellular automaton were introduced very early in the study of excitable media in the work of Wiener, and Rosenblueth [33]. Very early, since the use of computers was minimal at that time (in 1946, the date of publication), in such a way that the authors did not implement the cellular automaton in any computer, but used them in a purely theoretical manner. Despite this, they were able to explain how reentry waves circulate through obstacles. Subsequently, the use of cellular automaton to model excitable tissues became an area of continuous development, [1], [6].

The model we use can be inscribed in the line of the model introduced by Greenberg-Hastings [13], but with only one excited state and three refractory states, corresponding respectively to the Zhang and Holden states [35]: state 1 to the exciting state, 2, 3 to the excited state, and 4 to refractory state. At each discrete time *t*, the corresponding state *V_ij_*(*t*), where the subscript *i, j* is related to the cell’s position within a rectangular array. The update rule for the state of each cell is the following:

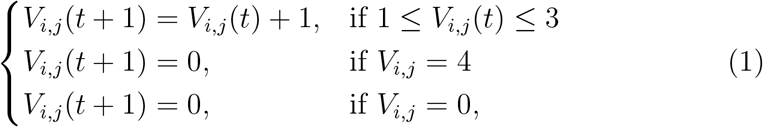

unless *V_i+1,j_* = 1 or *V_i-1,j_* = 1 or *V_i,j+1_* = 1 or *V_i,j−1_* = 1 in which case *V_i,j_*(*t* + 1) = 1.

According to the author’s experience, four states different from zero is the minimum number of states to produce *in silico* self-reproducing waves with the highly diffusive rules in (1). Note that once a cell starts to be excited it passes through the states sequence 1, 2, 3, 4, 0, without changing, which warranties uniqueness of the waves like that provided by the uniqueness of solutions in ODE. We used in a first step a rectangular arrangement of cells, but more complex distributions are studied latter in this paper. In [6] uniformly distributed arrangement of cells is studied as well as uniformly distributed weight for a cell’s influence on its neighbors. For automaton we keep the simplest model so that the threshold number for each cell is 1 in any one of its neighbors. So that as a main difference of the author’s paper with [6] is that to produce complex behavior of a net of cells such as fluttering is not required random distribution of neighboring cells not random threshold for activation. These models are useful because they have four stages as ordinary differential equations models do, even though, they are very elementary.

#### 2.1.2. Ordinary differential equations models

For this part, although only the experiments with the Barkley [5] model are reported, the Fitzhugh-Nagumo model and the Aliev-Panfilov model whose description is elsewhere [11], [22], [2] were also used. Since results obtained for flutter, and fibrillation are similar to those obtained with the Barkley model, Fitzhugh-Nagumo, Aliev-Panfilov models are not reported.

If a suitable geometry of the cell system is introduced (see section 2.2), it is possible to represent fibrillation and flutter phenomena with all these models. They are two-variable models, as is well known, and they are dynamical bi-stable systems. For these systems, the existence of limit cycles is well established in the mathematical theory, and even *analytical* approximations of physiologically relevant limit cycles in a region between heteroclinic trajectories are possible to calculate [17]. The main difference among them is related to better fitness to experimental data of action potential curves and other physiological parameters such as cycle length, et cetera, which the reader may find in [2].

The Barkley model used in this paper is the following

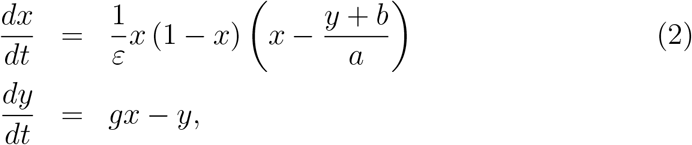

where *a, b, g, ε* are fixed parameters. Figure 1 shows an AP of Barkley model with initial conditions *x*(0) = 0.4, *y*(0) = 0,

As cellular automaton do, all ordinary differential equations models have four states. A rest state corresponding to a minimum value of the AP variable; an exciting state, which corresponds to a negative derivative of the AP profile; an excited state, which corresponds to the maximum of the AP profile; and a refractory state, which corresponds to a positive derivative of the AP profile. In figure 1 (b) the rest state is represented in the deepest blue color, and excited state is represented in the darkest red. Dynamical systems, even of two variables, represent more accurately actual excitable cells than automaton, but they are, of course, more costly computationally.

For cellular automaton we used a standard method of giving to each of the four states one color and plot the matrix in a rectangular arrangement corresponding to each time. For the ordinary differential equations models to solve the systems we use the Runge-Kutta method of order four with absolute error tolerances of 10^-6^. Observe that in any of our study cases, i. e., with automaton or with ODE systems a wave front can only propagate into a region in the resting state.

### 2.2. Cell-to-cell Nets Geometry

When using cell-to-cell models, the geometry in cellular systems can be determined by considering the geometry of the individual cells and how they are connected. For example, the working cells in the auricula in the heart are mostly cylindrical and are connected in a way that favors the longitudinal transmission of Action Potentials [30]. By comparison, brain cells have extremely branched forms, and their connections can reach a complexity that is far from being understood in its entirety [27]. In the following system,

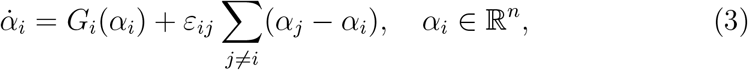

where *n* represents the number of variables of each cell. The geometry of the individual cells is intrinsically included in the mathematical model 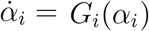, while the geometry of the network (and hence the diffusivity) is determined by the values of *ε_ij_* different from zero. So, equation (3) can be written as a vectorial equation with *α* = (*α*_1_,…, *α_n_*), *G*(*α*) = (*G*_1_(*α*),…, *G_n_*(*α*)). So, for two variables with the Barkley model in equation (2), the system (3) has the form

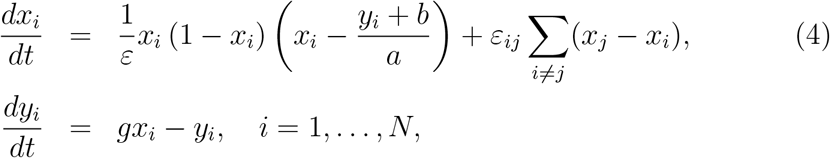

where *N* is the number of cells in the system. Notice that only the *x_i_* variables are coupled as corresponds to variables related to the Action Potential in the heart’s cells.

In practice, the use of rectangular and cubic matrices are the most commonly used [8], [12], [24], [36], without considering the complex geometry of the cytoarchitecture of the network that exists in the tissues of living beings. Systems made of systems of equations of the form (3) are known as *weakly connected networks* (WCN) and have a vast number of applications in neurophysiology [14]; cardiology [36], [8], [24]; and many other sciences. In general WCN have applications in every system of cells connected through gap junctions.

#### 2.2.1. Obstacles

There are two types of obstacles to be considered: dynamical, and static. Dynamical obstacles are formed by individual or groups of cells which are in a refractory or excitable or excited state. Static obstacles are formed by objects that do not change in time. They may correspond to fibroblast or adiposity in real tissue. In this article only dynamical obstacles are considered.

#### 2.2.2. Tiling

One of the central thesis of this work is that an intricate connection between cells is fundamental in the generation of fluttering and fibrillation. We take as a paradigm of cells connections, and hence of the intrinsic geometry of the cells, those of the working cells in the auricula and the ventricle in the heart. In the literature, histological studies of heart’s cells are available [30]. Nevertheless, mathematical models including real geometry of the cells are more scarce. Spach and Heidlage [31, Fig. 1] give a schematic representation myocardial architecture of 33 cells in a two-dimensional array. Following their representation, Figure 3 depicts a two dimensional model of cells, but our model is not based in real cells as in [31] but in a distribution generated by an aperiodic tiling called “table” which we describe below. After the connectivity of the cells is fixed it is possible to establish a correspondent cells geometry as in Figure 3. Note that the random-like distribution of the cells is not for real, in Figure 3 are represented the cells in the up and left corner in Figure 4. This distribution can be verified noting that the code of colors corresponds to the same cells, meaning green for cells with six connections, pink for cells with five connections, et cetera. Observe that in our figure, cells connections occur only in the vertical direction where most of the standard electrical coupling between cells have place [30].

**Figure (1).**
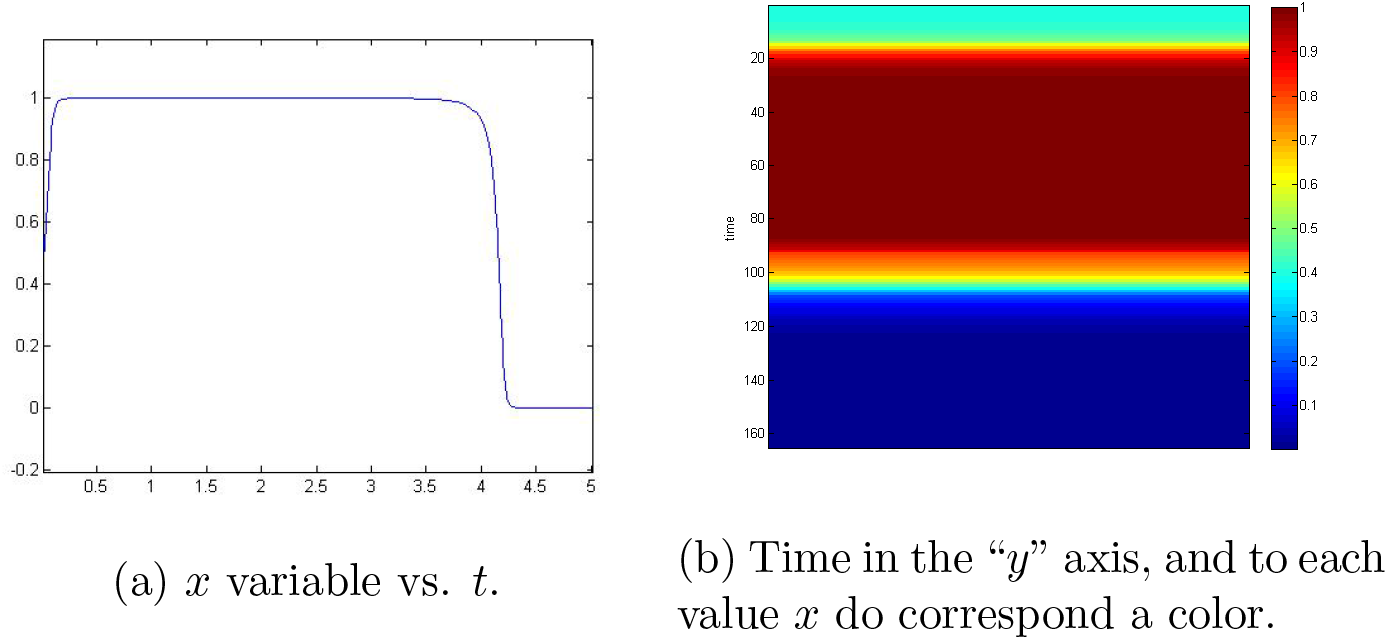
(a) Plot of variable *x* variable vs. *t* of equation (2). (b) To each value of *x* of the left plot corresponds a color to each height, the time is in the vertical axis. The colorbar corresponds to the states of Barkley model in the videos and plots included in this paper. Deep blue corresponds to rest state *x*(*t*) = 0. The darkest red corresponds to *x*(*t*) = 1.

**Figure (2).**
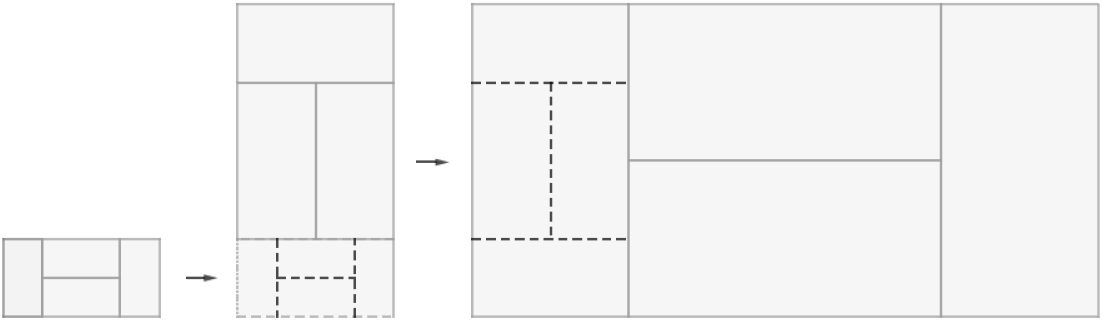
The procedure presented in the Figure is repeated several times to produce a “table”. Note the fractal structure obtained.

**Figure (3).**
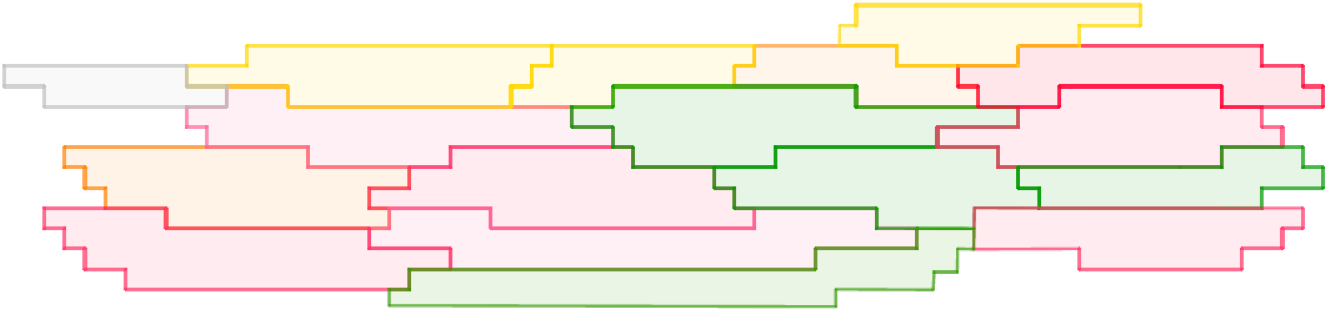
Here is shown one of many possible representations of working cells in the heart. They are not distributed randomly but following the *table* distribution described in Figure 4, where the reader can find the color’s code. The arrangement here presented corresponds to the up and left corner of Figure 4.

**Figure (4).**
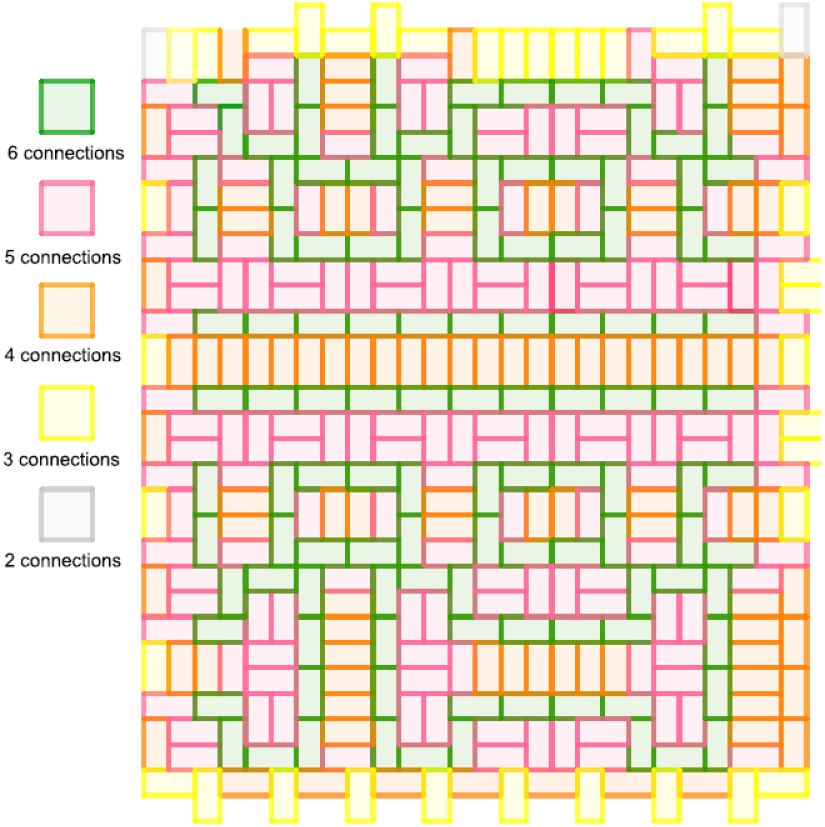
Example of a “table” tiling showing the number of connections of each cell. Cells in green are connected with six cells, cells in pink are connected with five cells, et cetera.

The values *ε_i,j_*· different from zero in equation (3) give a matrix of connections between cells which represents the geometry of the entire net. In this paper, we use a “ table “ distribution of connections among cells. A *table* is a polygon belonging to the class of rep-tiles that can be tiled by a finite number of smaller, congruent copies of itself (see [26], where the properties of “ table “ as tiling-dynamical-system are studied). We used this tiling for the following reasons. a) Is an aperiodic tiling of the plane so that some degree of complexity is intrinsic in the matrix of connections. b) The tiling is self-similar, so it does possess fractal structure. c) Each cell is connected to an average of 4.86 cells which is a good 2D approximation compared with an average of 9.1 reported from experimental measures by Hoyt et al. [15] for three-dimensional structures. Besides, the connectivity approaches that of the cells in the Spach diagrams of Figure 1 in [31], which represents a sample of two-dimensional tissue cells with an average two-dimensional connectivity of 4.66 cells. d) The “ table “ setting provides, in the author’s opinion, the simplest arrangement which satisfies the last mentioned properties.

Figure 4 shows a *table* used in our in silico experiments. This arrangement is only one sample of an infinite number of such aperiodic tilings. In order to set a matrix corresponding to the tiling in Figure 4 it is necessary to assign to each cell a number. Then, once the assignment is completed, the matrix of connections can be settled down. Algorithms to construct *table* and other aperiodic tiling are well known, but algorithms to set the number of connections in a given tiling and to write down the associated matrix of connections are not known, to the best of the author’s knowledge.

The recursive procedure to produce the self-similar “table” is shown in Figure 2.

### 2.3. Flutter, Fibrillation, and pseudo-ECG mathematical definitions

Although our interest is mainly focused on anisotropic excitable media in general and not just on the heart, the heart also seems to be the most natural medium for cell-to-cell modeling for apparent reasons. In this paper, we depart from the classical definitions of flutter and fibrillation given in the seminal work of Wiener and Rosenblueth [33]. Given that this article is old (1946) and somehow hard to find we quote:

“Fluter consists of a wave or waves of activity in a conducting system with a regular cyclic recurrence of paths, and therefore with a well-defined regular wave-front and period. Flutter differs from the ordinary heart-beat in that flutter activity is continuous, while beats are separated by periods of total rest (diastole of cardiac muscle). This definition agrees with both the clinical and experimental observations. Fibrillation consists of a continuous activity over randomly varying paths in a network of connected conducting elements.” Taking into account the last historical considerations the following definition applies to the rest of the article.

*Definition*. Flutter and fibrillation are reentrant waves of excitation which remain in a self-perpetuating steady state; flutter having a periodic (or nearly periodic) pseudo-electrocardiogram (pseudo-ECG), and fibrillation having a non-periodic pseudo-electrocardiogram. Alternatively, we call fibrillation a steady-state, self-perpetuating pattern of systems of cells without a defined front or back wave, such as the phenomenon shown in Figure 5.

**Figure (5).**
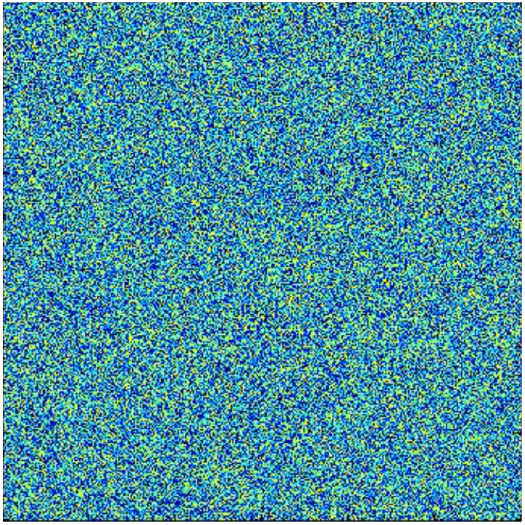
Cellular-automaton’s Fibrillation in a rectangle of 1 440 000 cells. This pattern is generated by increasing the number of cells with the same algorithm used to create Figure 8. The self-perpetuating, steady-state phenomenon is in video CA_fibrillation.avi

#### 2.3.1. Mathematical Pseudo-ECG

Following the classical definitions, we consider flutter and fibrillation as self-generating phenomena. Fluter is considering to be a periodic wave of waves contrary to fibrillation which is considered to be a highly complex nonperiodic wave or waves. A precise difference between flutter and fibrillation is provided by the pseudo-ECG which is calculated by the following formula:

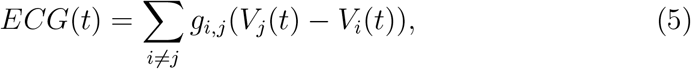

which is a non-weighted, two-dimensional version of the formula of Kazbanov et al. in [16]. Observe that the effect of the weights of formula (6) in [16] may be neglected since the number the cells that we considered is small (462). Hence the distances among them are small, and, more importantly, the geometry and thus the real distance between cells is not modeled in our study, so that weight due to the distance may not be considered.

## 3. Background

The simplest way to illustrate fluttering’s definition is by using a onedimensional system of cells in a closed ring. A ring is produced with a string of cells with periodic boundary conditions so that the first cell in the ring is connected with the last. As already mentioned, flutter must be a traveling wave self-perpetuating, re-entering, in a steady state. Without static fixed obstacles, system (1) produces a traveling wave which propagates in one dimension left and right. With a dynamic obstacle produced by one cell, one of the waves (right or left) cancels. A flutter appears with a minimum of five cells with our model of a cellular automaton. Figure 6 shows flutter with a minimum of five cells. As mentioned, the fifth cell is connected with the first, counting from left to right. In the subplot (a) the first cell is in refractory state 2, and the second cell is in state 1. As a result in subplot (b) a single wave propagates to the left. The first cell evolves to states 2,3,4, and 0 the rest state. In the subplot (c) appear (for the first time) the five possible states of a cell. Subsequent states from right to left are states 1,2,3, and 4, described in subsection 2.1.1. Flutter is evident in (e) when the first cell appears excited after being stimulated by the fifth cell as shown in (d). The complete sequence is in the video Cellular_Automata_annulus.avi. For Barkley ODE model given by equation (2) with *a* = 0.9, *b* = 0.5, *g* =1, *ε* = 0.02, eighteen cells are required to produce a flutter in an annulus. The complete sequence is in the video BARKLEY_annulus.avi.

**Figure (6).**
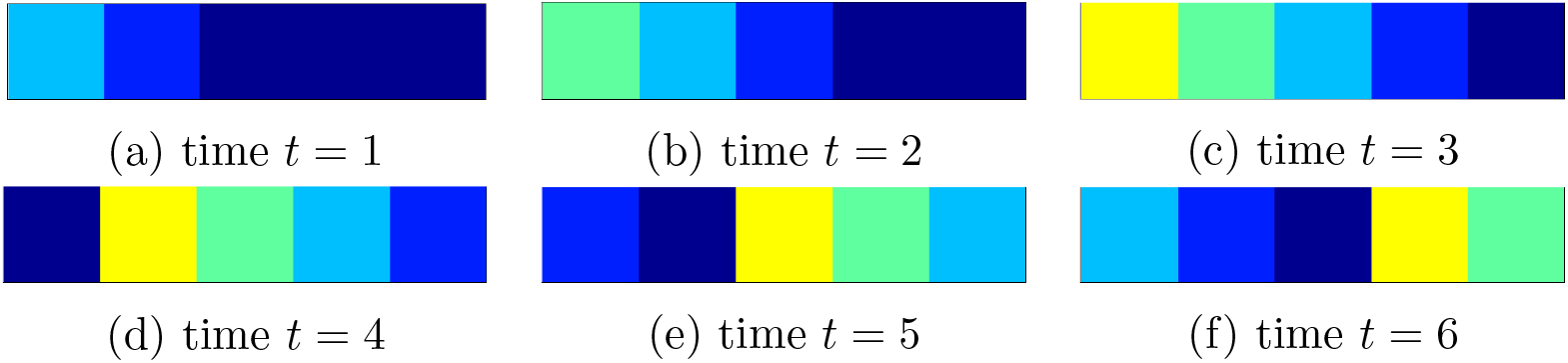
Sequence of activation in a ring of five cells with the cellular automaton of equation (1). The sequence is first from left to right and then, up to down. subplot (c) appear (for the first time) the five possible states of a cell. The extreme right cell in deep blue corresponds to state 0, corresponding to the rest state. Subsequent states from right to left are states 1,2,3, and 4, described in subsection 2.1.1.

Of course, there are different ways of producing a one-dimensional flutter. For instance, a first way, as described in [33], consists in stimulating one cell so that two waves emerge. After this, a second wave is produced by stimulating a cell in an immediate position to one of the waves’ back. Then the new wave collides with the old producing a single wave that runs forever in an annulus. The novelty of the cellular automaton model in this paper is that it is the minimal model with equation (1) and being minimal is essential in our following two-dimensional findings.

In two dimensions the mechanism for generating fluttering described by Wiener and Rosenblueth [33], consists in generating a wave which propagates radially in all directions. After a while, in the interior region of the wave, a second wave very close to the wave’s backis produced in such a way that the front of the second wave collides with the back of the first. The resulting wave after colliding with a static obstacle, such as vena cava, produces a self-generating wave.

In our automaton model to produce a flutter is not required an static obstacle. Instead we require a dynamical obstacle and the way of producing fluttering in one dimension is fundamental to produce the same cell-to-cell phenomenon in two dimensional diffusive media. We mean, by blocking some of the directions of propagation even with the minimum of one exciting cell. The author of this document found that blocking two directions of propagation for one cell with equation (1), patterns of fluttering emerge. In Figure 7 a fluttering emerges by blocking up and left cells of the cell in state 1. The complete sequence is in video Cellular_automata2D_b.avi.

**Figure (7).**
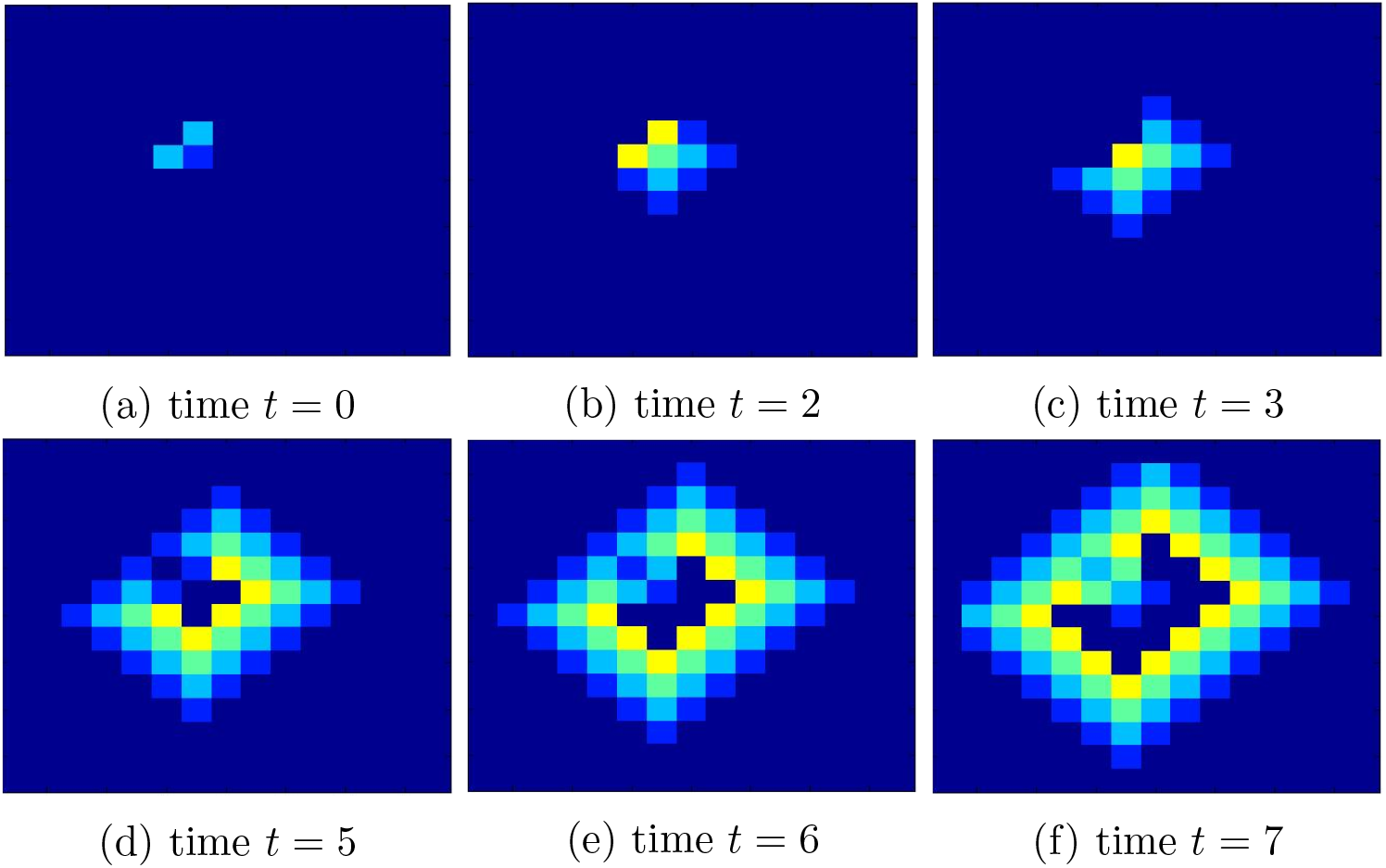
2D diffusion for cellular automaton. At *t* = 0 two cells en clear blue (state 2) block the cell in excitatory state. A periodic series of patterns emerge, which can be identified with which is called fluttering. At time *t* = 6 inside the wave can be observed the same initial configuration at *t* = 0. The complete sequence is in video Cellu-lar_automata2D_b.avi.

### 3.1. Mathematical background

The concept of observable has been used through the article; next, a mathematical definition is provided. Also, a comparison between PDE and cell-to-cell models is discussed.

#### 3.1.1. Observables

In mathematical modeling, the dynamics of the cells that form living beings (or that represent other excitable means) are represented by dynamic systems of form.

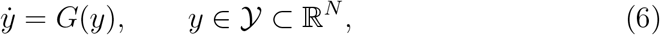

where the dot denotes the derivative with respect to time. So, cells are thought of as not wholly known (so far) dynamical systems, let us say

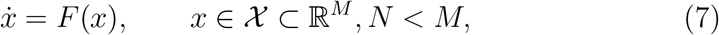

of which (6) is a representation, and scientists expect that the model *G* in some sense approaches *F*, which remain partially unknown. More formally, (6) is a model of (7) if there exists a continuous function (called observation [14]) 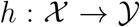 such that if *x(t)* is a solution of (7), then *y(t)* = *h*(*x*(*t*)) is a solution of (6). In practice, many information of system (7) is unknown, for instance, the dimension of the space (i. e., in this instance, the actual number of variables of the system). In many cases, a model could be a rough representation of the real system. As mentioned by Hoppenstead and Izhikevich [14], for example when (7) has a periodic solution and the model (6) is one dimensional, the observation *h*(*x*(*t*)) cannot be a solution of (6) unless h maps the limit cycle to a point. Actually, the existence of the function *h* and its properties are purely theoretic but allow us to speak about the relations of the real system (7) with the model in a mathematical fashion. As an example the variable *y*(*t*) = *h*(*x*(*t*)) is called an *observable*. In this article, the dimension of 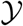 is bigger or equal than 2 so that we call *observable* to each of the component functions of *y*.

As science advances, mathematical models of cells include an increasing number of observables and each observable with an increasing refinement following experimental data. In this way, we obtain systems of complex differential equations that include an increasing number of equations. A typical example of this phenomena is the development in the study of the cells in the sinoatrial node in the heart (SAN) [7], [23], (for a detailed review of the SAN mathematical models see [19]).

However, this is just the first step on the way to modeling the actual cell tissue. A second step consists of forming a system of systems of equations by coupling variables among different systems, let say *n* different systems like the following

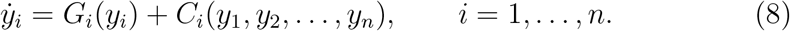

A very used example of a coupling functions *C_i_* are linear functions of the form 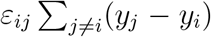 where *ε_ij_* are small (experimentally obtained) parameters and the values assigned to *j* depend on the geometry of the net. There are at least two ways of modeling excitable media. One is by utilizing partial differential equations to represent the diffusive nature of the media. Another is by establishing a system of cells, each cell, in turn, is a cellular automaton or is a system of ordinary differential equations (EDO). In the system of ordinary differential equations diffusion of the excitatory wave is modeled by coupling appropriate variables, for instance, the Action Potential (AP) in excitable biological cells. In this paper, we call *continuous* mathematical modeling to the first form, and the last form is what we call *cell-to-cell modeling*.

#### 3.1.2. Continuous vs. cell-to-cell modeling

Continuous mathematical modeling of anisotropic media such as ventricular tissue normally includes fiber patterns and the continuous rotation of the fiber axis [10], so that the equations have the form:

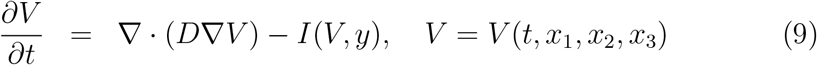

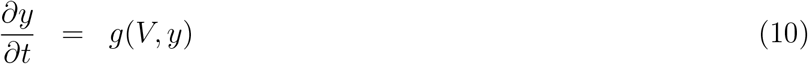

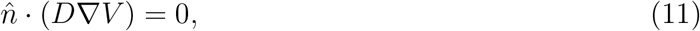

Where *V* is the membrane potential, *I* is the total current through the membrane, *y* is a vector of gate variables describing the dynamics of the various currents that constitute *I*, ∇*V* denotes the gradient operator, and *D* is a conductivity tensor divided cell surface to volume ratio times the membrane capacitance of the cell. We will show that a system of equations (4) is equivalent to the system (9), (10). Note that equation (11) represents Neumann boundary conditions. To begin with, observe that the tensor *D* is of the form

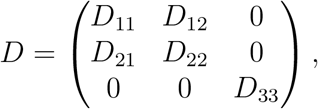

where *D_ij_* are functions of diffusivities parallel and perpendicular to the fiber, and *θ*(*x*_3_), the angle between the fiber to the axis of each plane. In the setting of [10], is easily shown that for a two-dimensional model, since *θ*(*x*_3_) ≡ 0, then *D* becomes

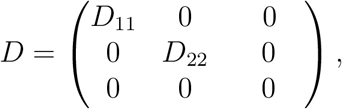

where now *D*_11_, *D*_22_ are constants, so that the elliptic operator ∇ · (*D*∇*V*) in equation (9) becomes simply

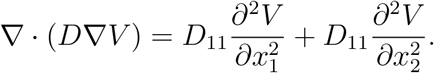

After discretization of the second partial derivatives we obtain

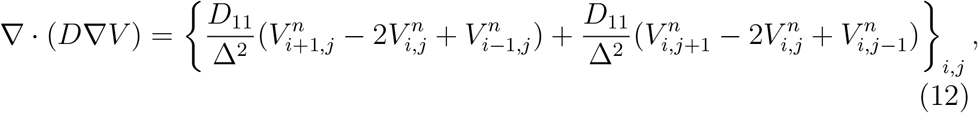

a formula which is valid for interior cells in the grid. Since in many articles including [10] the grid spacing is about Δ ≈ 200 — 300 *μ*m which is bigger than the length of the cell, ≈ 80 *μ*m [30], is worth to mention that PDE continuous approach is, in this case, not better than cell-to-cell modeling whatsoever. Moreover, note that equation (12) corresponds to a rectangular grid of square cells (of much bigger dimensions than actual heart cells), meaning a chessboard-like geometry of the cells. In this way a complex geometry such as that depicted in Figure 3 cannot be represented by the elliptic operator in equation (9) since such arrangement can not at all be represented with tridiagonal matrices such as those in equation (12).

A further consideration regarding the mathematics in this article must be considered. Flutter and arrhythmia will appear as solutions of systems of ODEs for a particular set of initial conditions, but somehow the solutions appear in some fashion unpredictable since they occur after global bifurcations of the parameters given by the conductivity and of the distribution of conductivity. Hence, only after integration of the systems will emerge more of the most striking patterns of the next section in an unexpected form. Although the spirals formed by Barkley model [3], [4], [5] and other models [9] have been extensively studied, the combination of spirals, collisions of spirals, and spirals emerging after a massive dynamic blocking leading to fibrillation are still of interest.

## 4. Results

In this paper, the waves mentioned in the definitions above are traveling waves with a defined front and back given by equation (1) (for the automaton case) or given by the ordinary differential equations of the different bistable ODE systems. As mentioned in section 2, systems with no apparent wave’s fronts and backs are noticeable. In this case, we call them fibrillation too, such as the phenomena depicted in Figure 8. Nevertheless, a periodic pseudo-ECG may appear after time in some of the systems formed with a small number of cells. While flutter is produced by the collision of traveling waves with dynamical or static obstacles at a macroscopic level as in the classic definition, there is a difference in this document with the standard definitions since microscopic (cell-to-cell) collisions are considered, and very intricate patterns may happen.

**Figure (8).**
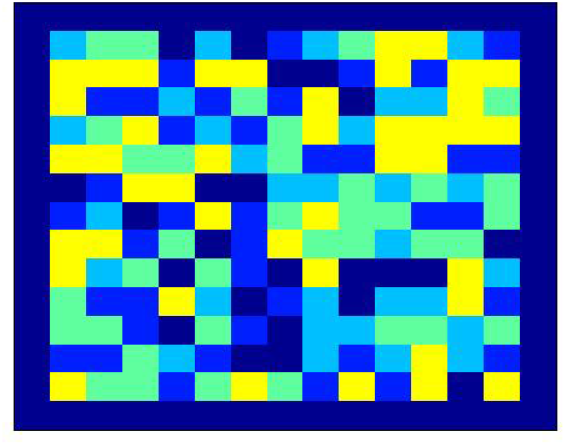
One of an infinite number of self-perpetuating patterns obtained with initial conditions mixing exciting and refractory cells.

### 4.1. Fibrillation at a microscopic level

The point here is that, in reality, in two dimensions an infinite number of self-perpetuating patterns will emerge by combining dynamic blocking with exciting cells at t=0, in contrast with the one-dimensional case. Figure 8 shows one of the infinite possible patterns where the blocking and excitation cells are setting at *t* = 0. The initial conditions correspond to a uniform distribution. Surprisingly, a self-perpetuating pattern appears in almost any initial random configuration. Surprisingly, since one can expect self-perpetuating patterns to be an unusual phenomenon, but it is not so with dynamic blocking. The video corresponding to Figure 8 is Random.avi.

After obtaining an infinite number of self-perpetuating patterns in cell-to-cell modeling, we are in the possibility of generating a model of automaton-made fibrillation by merely combining a significant number of the complex patterns obtained, and by introducing a bigger the number of cells. Figure 5 shows an square of 1200 × 1200 cells. Indeed, the phenomenon depicted in Figure 5 and video CellularAutomata_fibrillation.avi, corresponds more properly with fibrillation that with fluttering, given that not single waves with defined front and back are recognizable. The reader may ask if such dramatic mechanism shown in Figure 5 also occurs with more complex models than cellular automaton. Figure 14, shows a massive blocking for Barkley model cells which after a while becomes fluttering.

Fluttering, as the reader may recall, is associated with a well definite macroscopic steady-state wave which often present spirals as one depicted in Figure 9. Of course, since cells in the cellular automaton model are rectangles of 100 micrometers in order to bee visible macroscopically the wave length should be larger than the wave shown. In Figure 9, the wave length is only of four cells. In Figure 9, the wave length is only of four cells. By augmenting the number of refractory states in the equation 1 is possible to increase easily the wavelength of cellular automaton, but in this paper we are interesting in a setting of a minimum of parameters.

**Figure (9).**
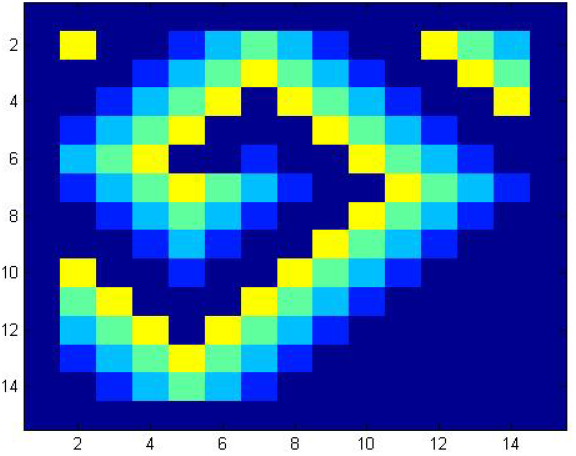
A flutter in spiral form with cellular automaton.

#### 4.1.1. Two dimensional fluttering and fibrillation with ODE

All the phenomena studied in subsection 4.1 are reproduced by using ODE, in particular by using Barkley model with equation (2). In cell-to-cell coupling models with ODE, fluttering is simulated, as in the case of cellular automaton, by blocking an exciting cell with cells in a refractory state. Fluttering is also obtained by blocking only one cell with two cells or by a more significant number of cells connected with the boundary and hence producing spirals. Figure 10 shows a spiral created with Barkley model with a “table” distribution for the connections between cells.

**Figure (10).**
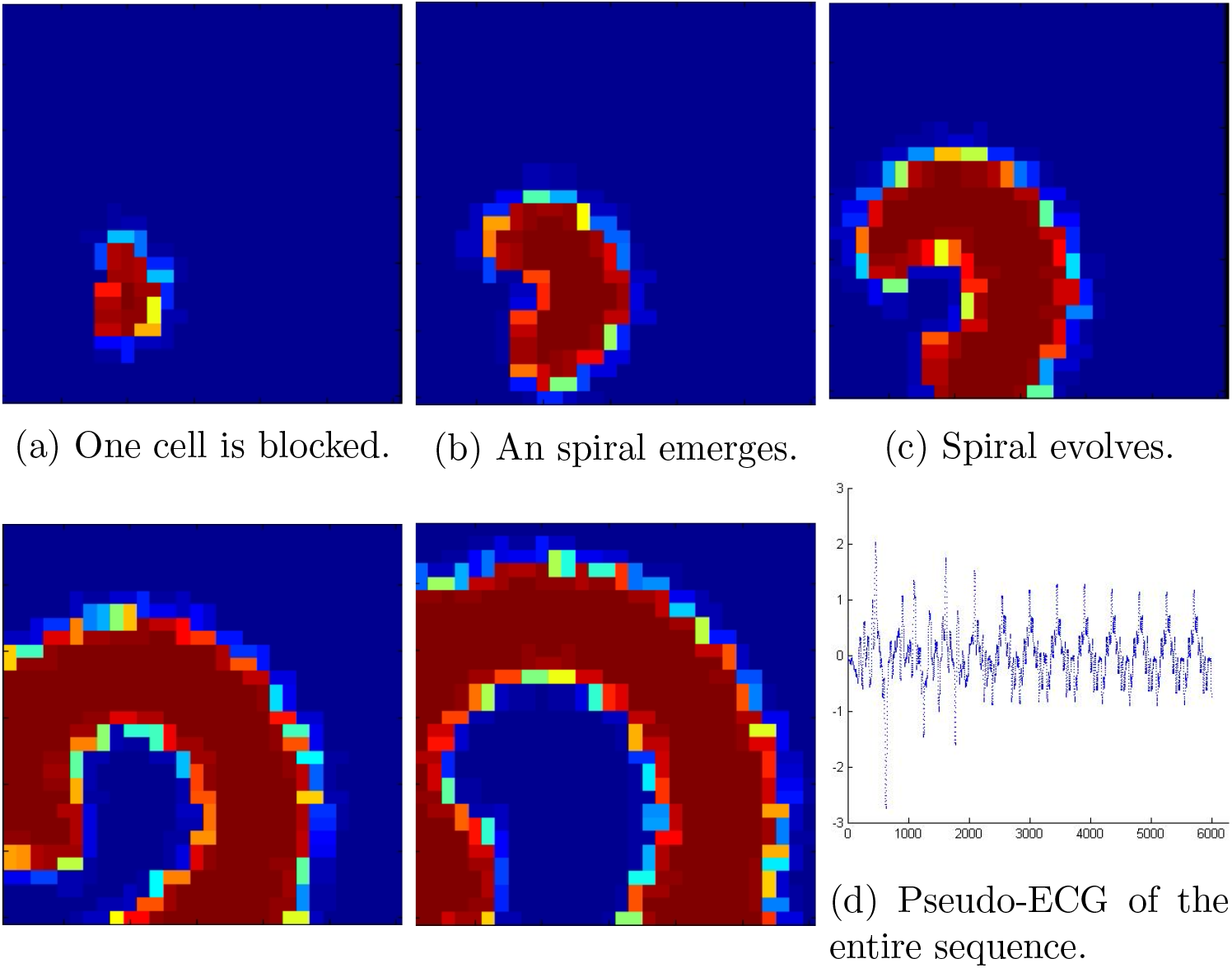
A typical fluttering spiral is produced by blocking one cell and normal conductivity. The sequence is from left to right, and up to bottom. The pseudo-ECG associated to the figure is in Figure 11.

Note that the pseudo-ECG Figure 11 prasents in the beginning an irregular non periodic behavior but after time *t* = 2000, the ECG becomes periodic, just as expected during fluttering.

**Figure (11).**
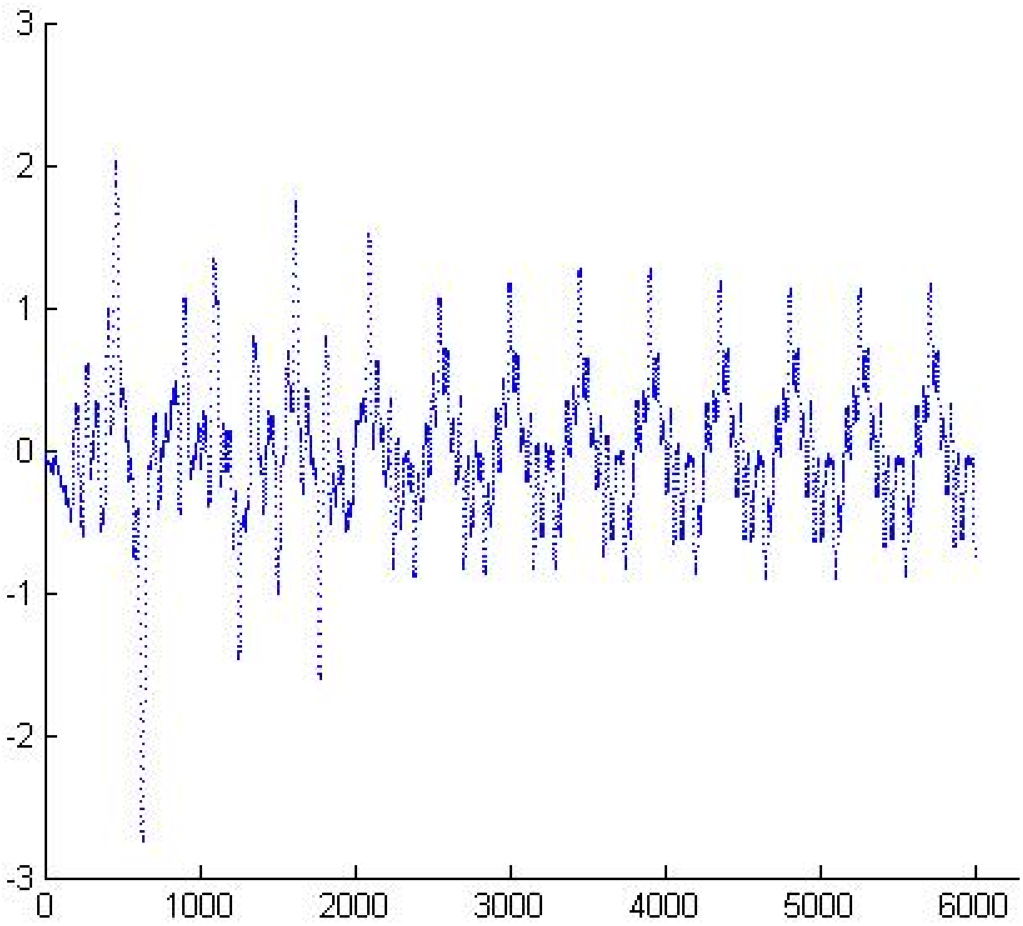
A flutter in spiral form with Barkley model in a system of cells with the distribution shown in figure 10. The pseudo-ECG presented here depicts periodic behavior after t = 2000, when the spiral becomes stable.

More interesting, as we present in this section, is to obtain models of the phenomenon of fibrillation without introducing a random variation of some parameters, which is the most common [6], [31]. Instead of using random connectivity in a random number of neighbors, the model of this paper uses the connectivity shown in Figure 4.

For the Barkley model given by equation (4) with *a* = 0.9, *b* = 0.5, *g* =1, the critical value is *ε_ij_* = 0.9 with the “table” distribution of figure 4. With the mentioned critical value given, the variable connectivity of each cell a variable resistance for diffusion causes a complex pattern which may be identified with local cell-to-cell fibrillation. That is, variable connectivity with a critical diffusivity value produces dynamic obstacles that induce fibrillation in the entire system. Hence, in Figure 12 is simulated a “more subtle” fibrillation than that modeled with the cellular automaton due to a different kind of dynamical obstacles given by critical connectivity combined with complex connectivity of each cell. The complete sequence of Figure 12 is shown in video 121blk105a110_462Kp9.avi. The pseudo-ECG associated with the fibrillation of Figure 12 is shown in Figure 13.

**Figure (12).**
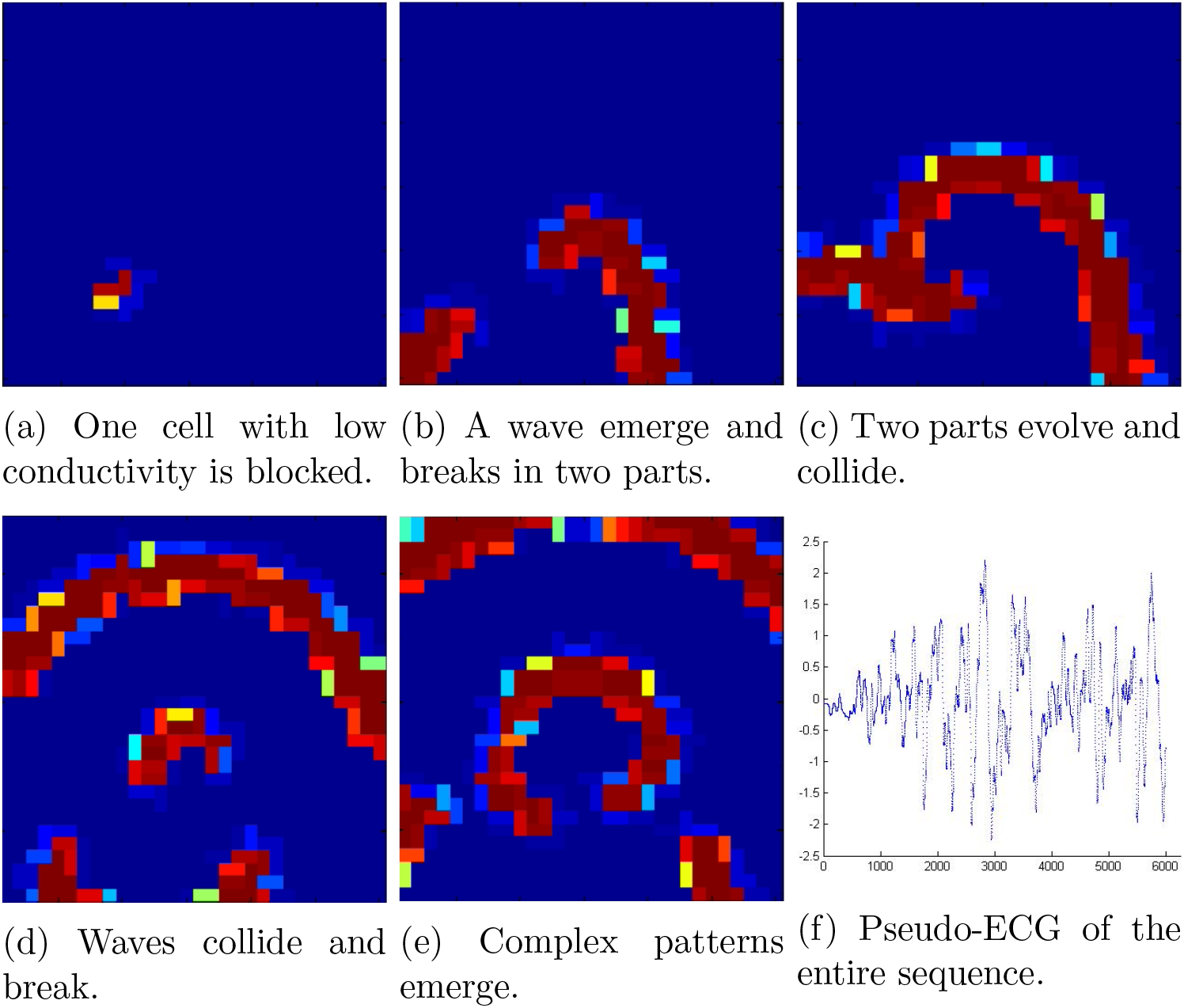
A single exciting cell without refractory blocking cells produces this sequence. When the net has critical connectivity, an intricate pattern of diffusion emerges due to the variable connectivity of the cells and low conductivity. Note how spirals broke into scrolls. The complete sequence is in video 121blk105a110_462Kp9.avi.

**Figure (13).**
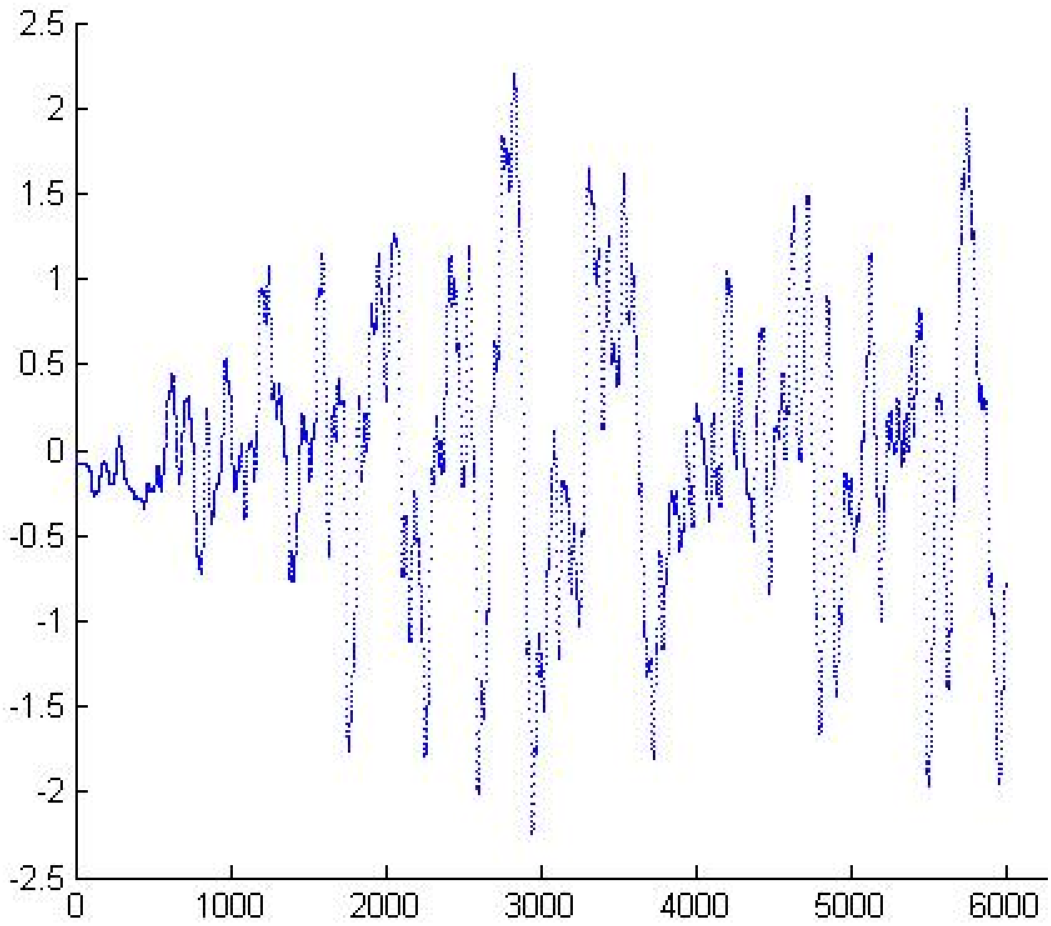
This pseudo-ECG shows the complex and aperiodic nature of the fibrillation depicted in Figure 12

#### 4.1.2. Fibrillation becomes fluttering

The last phenomenon presented here in Figure 14 is the spontaneous initiation of fluttering right after fibrillation produced by a massive blocking distributed uniformly through the net. In the heart, this mechanism is common shortly after open heart surgery (see [32] and references therein). The pseudo-ECG corresponding to Figure 14 is in Figure 15.

**Figure (14).**
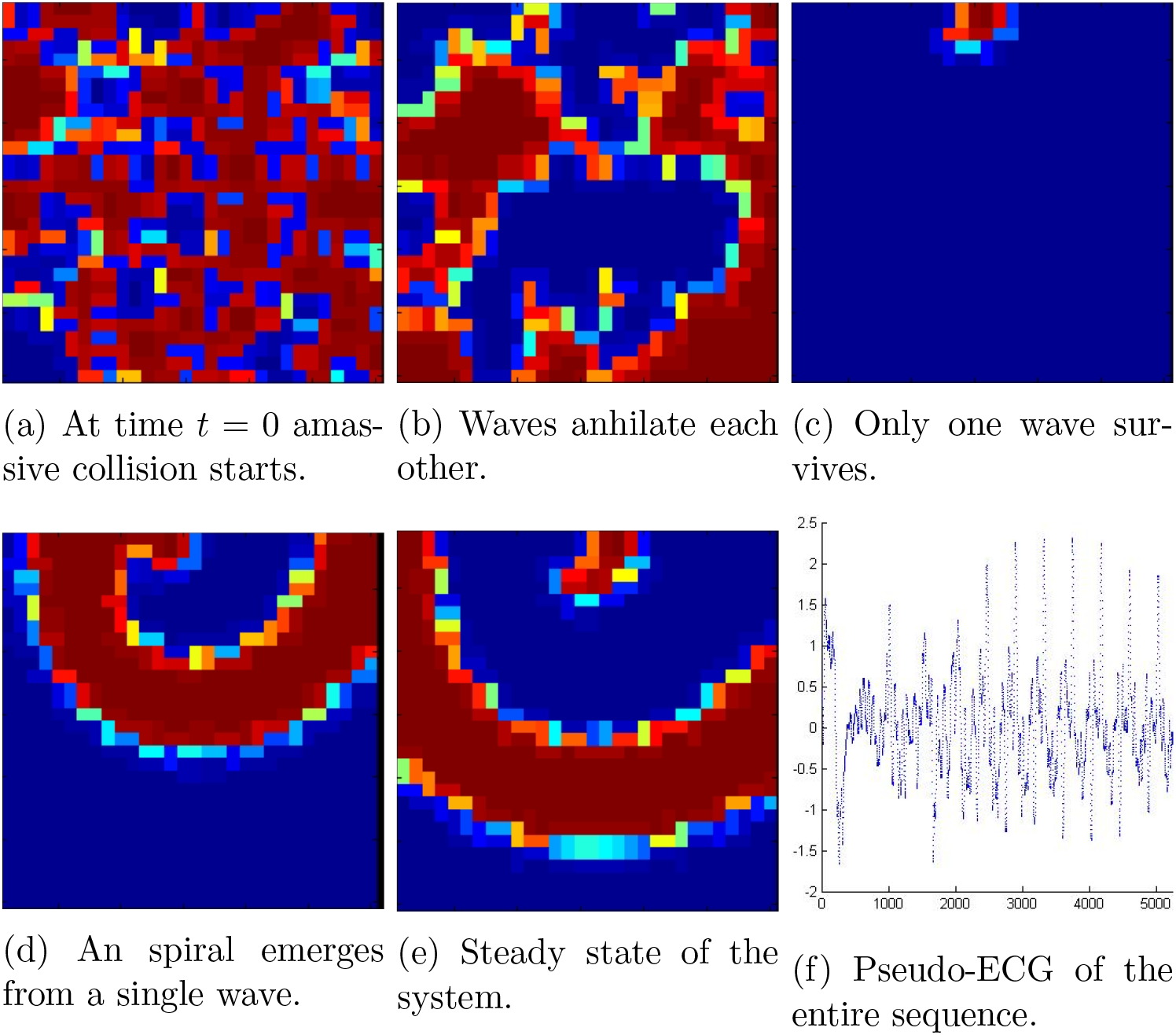
Massive blocking produces fibrillation which becomes fluttering. The complete sequence is in video BARKLEYtablaFIBR_FLUTT_1.avi. The pseudo-ECG 14f corresponds to Figure to the complete sequence. Observe how the complex activity from time *t* = 0 to *t* = 2500 may be associated with fibrillation. After *t* = 2500, the activity becomes nearly periodic corresponding to flutter phenomenon.

**Figure (15).**
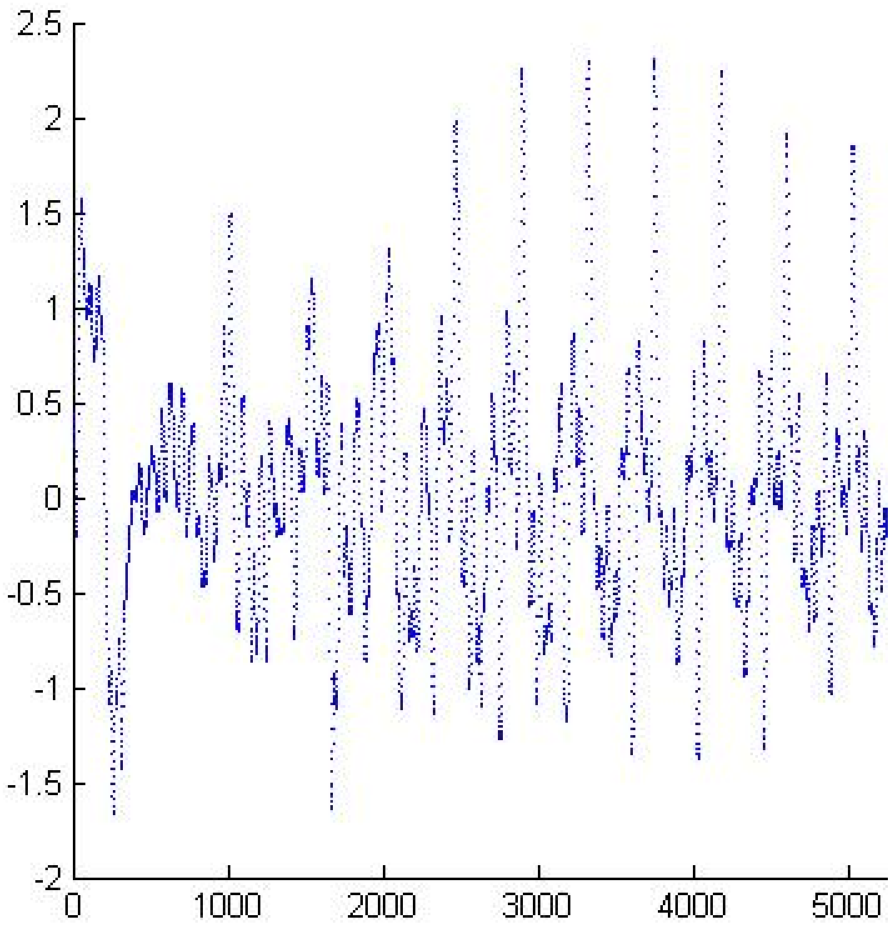
This pseudo-ECG corresponds to Figure 14. Observe how the complex activity from time *t* = 0 to *t* = 2500 may be associated with fibrillation. After *t* = 2500, the activity becomes nearly periodic corresponding to flutter phenomenon.

## 5. Discussion

In many articles, one and two-dimensional arrangements (for instance in [8] and references therein and [36]) are considered disregarding that the actual geometry of tissue is three dimensional. This reduction in modeling is a generalized attitude that can be understood under the dynamic of building models that go from the simple to the more complicated. Nevertheless, as Fenton et al., claim [9] in models of cardiac electrical activity “simulations in 3D have shown that the existence of purely three-dimensional breakup mechanisms.” Hence in order to obtain more accurate models of arrhythmias, it is necessary to work in 3D frameworks, but utilizing 2D layers make sense for the following reasons. In modeling auricular heart tissue it is known [10] that rotational anisotropy of fibers of ventricular muscle can be model by superposing and rotating two-dimensional layers of cells. To this aim, two ways to incorporate connectivity parameters in the cells are available: experimental histological data or stochastic or complex connectivity provided by mathematical models ad hoc. In this article, we used a “ rep-tile” model which is possible to extend to 3D nets of cells. Anyhow, the extension to 3D of the “table” model presented here is not straightforward, since the algorithms to create a matrix of connections corresponding to a table or other aperiodic, fractal arrangements are not available yet (this is part of a work in progress), although algorithms to produce many of such patterns are known (see for instance Rangel-Mondragon article [25]).

In this document, cell models with a high number of observables were not used (for definition, see section 3.1.1), since, as we illustrate in the section 4, the diffusivity provided, neither by PDE nor by cell-to-cell coupling models, requires complex dynamics in individual cells to produce fibrillation and flutter phenomena to the extent of the complexity studied in this document. More sophisticated models of individual cells are considered in work in progress where the influence of the nervous system for modeling is required.

Although simplified models in one or two dimensions may reflect certain experimental data [12], it may be that such simplifications do not represent the actual behavior of big groups of cells, for instance, in the transmission of the action potential of the system formed by mixing pacemaker and atrial cells. For example, in [20] it is shown that for specific mathematical models of atrial and sinus cells, the activation of the complex depends on the number of cells involved, as well as on the geometric distribution of the cells in the network. Taking this into account, modeling diffusive media with cells including a big number of observables may not be a trivial task, since even the stability of the numerical methods involved may be in chalenge. Moreover, since the real geometry of the cells must be taken into account, given the results obtained in silico in this document and elsewhere results, real data of local topology of diffusive tissue must be incorporated, when available, in order to model big groups of cells. As mentioned in section 4 our representation of groups of cells in figure 3 may not correspond to any real biological tissue, but only constitute the first approach of statistical data of a mean of the number of observed connections in the atrial heart tissue. A refinement of this data is required for the two-dimensional layers forming atrial and ventricular tissue in the heart.

On the other hand, many interesting works such as [31] study the stochastic distribution of inhomogeneities at the cellular level that can cause cardiac propagation to be stochastic. In contrast, in this article without considering the stochastic setting, we obtained a complex propagation in the diffusive tissue, despite the simplicity of the included models, only by varying the local topology of the network. Nevertheless, variable conductance may be included in future work either employing stochastic distributions or, as was done in this article, introducing some aperiodic pattern of the distributions of the different conductances referred in [31] of the individual cell membranes.

## 6. Conclusions

In modeling cell-to-cell in this document, we found that a very complex self-perpetuating diffusion patterns arise utilizing a massive blocking of cells in exciting state. This complexity emerges even in utilizing very simple chessboard-like distribution of the simplest automaton that generates without a massive blocking a solitary wave. One remarkable property of the cellular automaton nets of this document is that reentrant waves are formed in a vast variety of initial conditions contradicting the intuitive folk thinking that arrhythmia phenomena are exceptional in diffusive media, especially in considering fibrillation.

After the cellular automaton exploration and taking the results obtained as a motivation, we continue studying arrhythmias with ODE systems models, but on more complex arrangements than chess-board topology nets. We introduce a net with “table” rep-tile distribution in which fibrillation, fluttering, and a sequence fluttering-fibrillation phenomena like emerge. In this way, the two basic types of arrhythmia were modeled in two-dimensional tissue with a degree of complexity given by the non-periodic, fractal distribution connections in the “table”. The interesting fact is that it is possible to model a complex-like fibrillation phenomenon by introducing a certain degree of complexity in the distribution of neighbor cells (for instance with a “table”), instead of using random distribution. To the best of the knowledge of the author of this paper, this is a novelty. Moreover, the author found by integrating the existence of a critical value of conductivity among the cells, such critical value emerges with a matrix of connections given by the arrangement in figure 4. Low conductivity of cells in the heart is associated with ischemia [18], and in experiments, conductivity may be lowered pharmacologically by heptanol [6]. In this way, modeling fluttering by lowing conductivity in our model of simple two-variable ODE model by adding simply certain complexity in the distribution of cells could be of some relevance in mathematical modeling and in computational simulation.

With cellular automaton the wave length is of four cells, for Barkley model the wave length varies according with the connectivity parameter. Wave length is relevant to produce a complex steady state like the depicted in Figure 5. Such a complex pattern is not produced with Barkley model even with critical conductivity parameter. Instead of that static-like phenomenon in some in silico experiments emerge a number of self-perpetuating waves which collide, leading to a complex pseudo-ECG which anyhow may be identified with fibrillation of the system. An example of this arrhythmia is shown in Figure 16.

**Figure (16).**
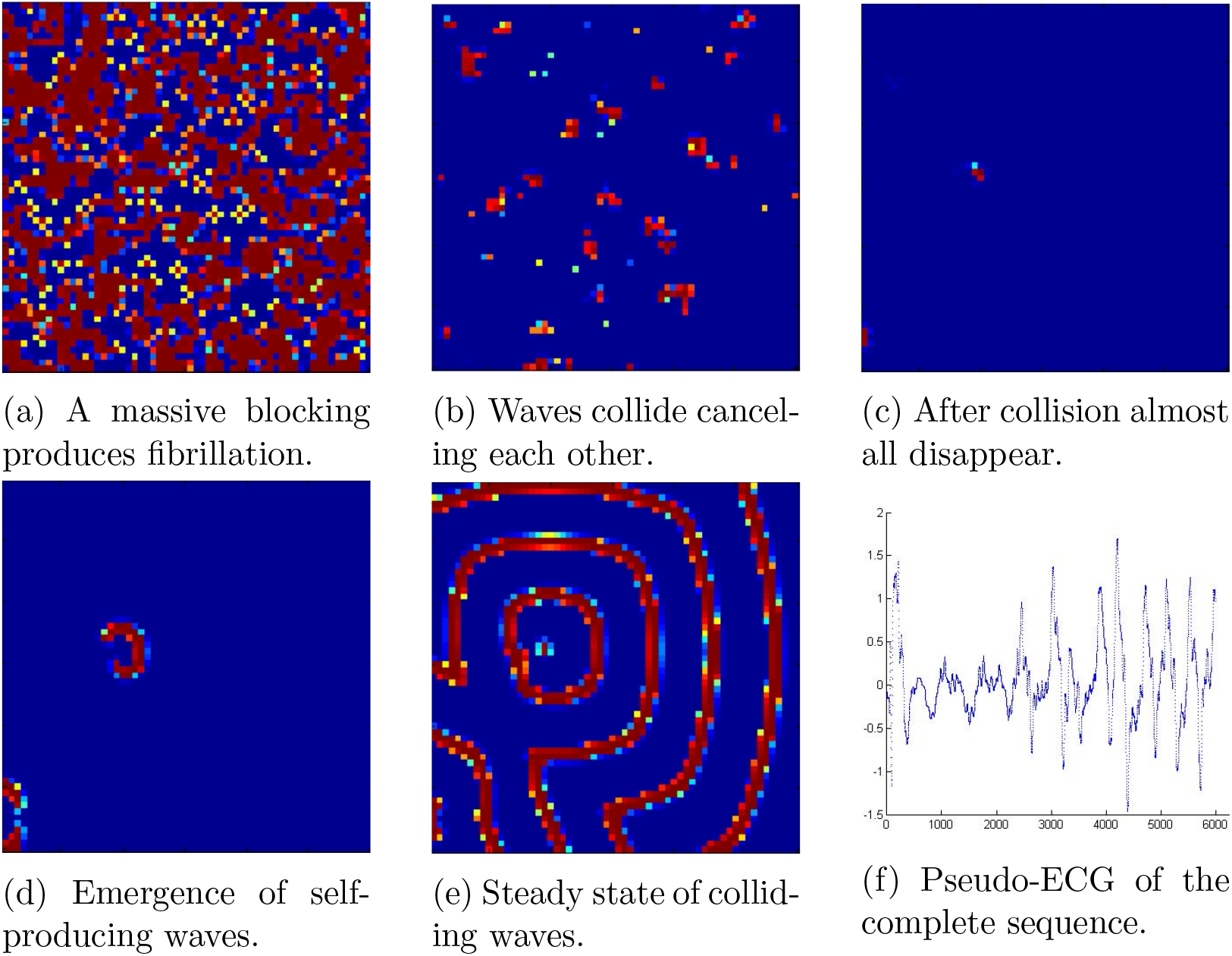
Barkley model after massive blocking of individual cells with low conductivity produces a complex pattern that according with the pseudo-ECG 16f can be identified with fibrillation. Here we show a system of 60×60 cells.

## References

[1] Adamatzky A., Martínez G. J. Editors Designing Beauty: The Art of Cellular Automata. ISSN 2194-7287 ISSN 2194-7295 (electronic) Emergence, Complexity and Computation ISBN 978-3-319-27269-6 ISBN 9783-319-27270-2 (eBook) DOI 10.1007/978-3-319-27270-2

[2] Aliev R. R., Panfilov A. V., A Simple Two-variable Model of Cardiac Excitation. Chaos, Solitons & Fractals Vol 7, No. 3, pp 293–301. 1996

[3] Barkley D., Kevrekidis, I. G., A dynamical systems approach to spiral wave dynamics. Chaos 4 (3), 1994.

[4] Barkley D., Euclidean Symmetry and Dynamics of Rotating Spiral Waves. Physical Review Letters V 72, No. 1., 3 January 1994.

[5] Barkley D., et al. Spiral-wave dynamics in a simple model of excitable media: The transition from simple to compound rotation. Physical Review A, Rapid Communications, vol. 42, No. 4 15 August 1990.

[6] Bub G., et al. Spiral Wave Generation in Heterogeneous Excitable Media. Physical Review Letters Vol 88, n. 5 February 2002.

[7] Castellanos P., Godinez R., Autonomic nervous system regulation of the sinoatrial cell depolarization rate: Unifying computational models. 37th Annual International Conference of the IEEE Engineering in Medicine and Biology Society (EMBC), pp 43–46, Aug 2015.

[8] Garny A.,Kohl P.,Hunter P. J., Boyett MR, Noble D., One-Dimensional rabbit sinoatrial node models. J Cardiovasc Electrophysiol. (2003) 14:S121S132. doi: 10.1046/j.1540.8167.90301.x

[9] Fenton F. H., et al. Multiple Mechanisms of spiral wave breakup in a model of cardiac electrical activity. Chaos, Vol. 12 No. 12, 2002.

[10] Fenton F., Karma A. Vortex Dynamics in three-dimensional continuous myocardium with fiber rotation: Filament instability and fibrillation. Chaos Vol. 8 No. 1 1998.

[11] FitzHugh R., Impulses and physiological states in theoretical models of nerve membrane. Biophys. J. 1, 445–465 (1961)

[12] Garny et al. Dimensionality in cardiac modelling. Progress in Biophysics and Molecular Biology 87 (2005) 47–66.

[13] Greenberg J., et al. Spatial Patterns for Discrete Models of Diffusion in Excitable Media. SIAM J. Appl. Math., 34(3), 515523.

[14] Hoppensteadt F. C., Izhikevich E. M., Weakly connected Neural Networks. ISBN 0-387-94948-8 Springer-Verlag New York Berlin Heidelberg SPIN 10557261.

[15] Hoyt R. H., Cohen M. L., Saffitz J. E. Distribution and three-dimensional structure of intercellular junctions in cannine miocardium. Circ Res. 1989; 64: 563–574.

[16] Kazbanov, I., V., et al. Effects of Heterogeneous Diffuse Fibrosis on Arrhythmia Dynamics and Dynamics. Nature Scientific Reports—20835— DOI: 10.1038/srep20835.

[17] Keener J., Sneyd J., Mathematical Physiology I: Cellular Physiology. Second Edition. Section 6.2 pp. 231–235 Springer, ISBN 978-0-38775846-6.

[18] Kleber A., G., et al. Electrical Uncoupling and Increase of Extracellular Resistance After Induction of Ischemia in Isolated, Arterially Perfused Rabbit Papillary Muscle. Circ Res. 1987 Aug;61(2):271–9.

[19] Li P., Lines G. T., Maleckar M. M., Tveito A., Mathematical models of cardiac pacemaking Function. Frontiers in Physics. October 2013, Vol. 1 Article 20.

[20] López G., et al. Cell-to-cell modelling of the interface between atrial and sinoatrial anisotropic heterogeneous nets. Computational Biology and Chemestry 68 (2017) 245–259.

[21] Lewis, T. The Mechanism and Graphic Registration of the Heart Beat. London 1925.

[22] Nagumo J. S., et al. An active pulse transmission line simulating nerve axon. Proc. IRE. 50, 2061–2071

[23] Noble D., Modification of the Hudgking-Huxley equations applicable to Purkinje fibre action and pace-maker potentials. J. Physiol. (1962), 160, pp.317–352.

[24] Oren R. V., Clancy C., E., Determinants of heterogeneity, Excitation and Conduction in the Sinoatrial Node: A Model Study. PLoS Comput Biol 6(12): e1001041. doi:10.1371/journal.pcbi.1001041.

[25] Rangel-Mondragon J., Polyominoes and Related Families. The Mathematica Journal 9:3 2005 Wolfram Media, Inc.

[26] Robinson E. A., On the table and the chair. Indag. Mathem.,N.S., 10 (4), 581–599.

[27] Rubinov M., Sporns O., Complex network measures of brain connectivity: Uses and interpretations. Neuroimage 52 (2010) 1059–1069.

[28] Saoudi N., et al. A classification of atrial flutter and regular atrial tachycardia according to electrophysiological mechanisms and anatomical bases. European Heart Journal (2001) 22, 11621182 doi:10.1053/euhj.2001.2658, available online at http://www.idealibrary.com

[29] Savalia S., Emamian V., Cardiac Arrhythmia Classification by MultiLayer Perceptron and Convolution Neural Networks. Bioengineering 2018, 5, 35; doi:10.3390/bioengineering5020035

[30] Shimada T. et al. Cytoarchitecture and Intercalated Disks of the Working Myocardium and the Conduction System in the Mammalian Heart. The anatomical Record Part A 280A:940–951 (2004).

[31] Spach M. S., Heidlage J. F., The stochastic Nature of Cardiac Propagation at a microscopic Level.

[32] Waldo A, L., Mechanisms of atrial flutter and atrial fibrillation: distinc entities or tow sides of a coin? Cardiovascular Research 54 (2002) 217–229.

[33] Wiener N., Rosenblueth A., The Mathematical Formulation of the Problem of Conduction of Impulses in a Network of Connected Excitable Elements, Specifically in Cardiac Muscle. Archivos del Instituto de Car-diologia de México;ao 16 Tomo XVI 1946 Nos. 3 y 4.

[34] Wiener N. and Wintner A. The discrete chaos. Amer. J. Math., 65: 279–298.

[35] Zhang H., Holden A. Chaotic Menander of Spiral Waves in the FitzHugh-Nagumo System. Chaos, Solitons & Fractals Vol 5, Nos 3/4, pp 661–670, 1995.

[36] Zhang H., Holden A., Kodama I., Honjo H, Lei M., Varghese T., et al. Mathematical models of action potentials in the periphery and center of the rabbit sinoatrial node. Am J Physiol Heart Circ Physiol. (2000) 279:H397H421. Available online at: http://ajpheart.physiology.org/content/279/1/H397

[37] Zacharia A. M., et al. Cardiac Arrhythmia Classification Using Atrial Activity Signal. Procedia Technology 24 (2016) 1406–1414.

